# Astrocyte ezrin defines resilience to stress-induced depressive behaviours in mice

**DOI:** 10.1101/2024.09.10.612240

**Authors:** Si-Si Lin, Bin Zhou, Si-Le Liu, Xing-Ying Ren, Jing Guo, Jing-Lin Tong, Bin-Jie Chen, Ruo-Tian Jiang, Alexey Semyanov, Chenju Yi, Jianqin Niu, Peter Illes, Baoman Li, Yong Tang, Alexei Verkhratsky

## Abstract

Astrocyte atrophy is the main histopathological hallmark of major depressive disorder (MDD) in humans and in animal models of depression. Here we demonstrated that manipulating with ezrin expression specifically in astrocytes significantly increases the resilience of mice to chronic unpredictable mild stress (CUMS). Overexpression of ezrin in astrocytes from prefrontal cortex (PFC) rescued depressive-like behaviours induced by CUMS, whereas down-regulation of ezrin in astrocytes from PFC increased mice susceptibility to CUMS and promoted depressive-like behaviours. These behavioural changes correlated with astrocytic morphology. Astrocytes from PFC of mice sensitive to CUMS demonstrated significant atrophy; similar atrophy was found in astrocytes from animals with down-regulated ezrin expression. To the contrary morphology remains unchanged astrocytes in animals resistant to CUMS and in animals with astrocytic overexpression of ezrin. Morphological changes also correlated with ezrin immunoreactivity which was low in mice with depressive-like behaviours and high in mice resistant to stress. We conclude that Ezrin-dependent morphological remodelling of astrocytes defines the sensitivity of mice to stress: high ezrin expression renders them stress resilient, whereas low ezrin expression promotes depressive-like behaviour in response to chronic stress.

## Introduction

Psychiatric diseases in general, and diseases of mood in particular, arise from an aberrant functionality of neuronal networks associated with pathological changes in neuronal excitability, synaptic transmission, synaptic plasticity, network architecture, and nervous tissue homeostasis ^1–5^. Uncompensated chronic stress is one of the key aetiological factors in major depressive disorder and stress-induced depressive behaviours; notably, stress affects structures in the prefrontal cortex (PFC) that is instrumental in stress processing ^6^. These structural changes reflect loss of tissue homeostasis and are manifested by synaptic loss and decrease in neural cells densities ^7^. Emergence of depressive symptoms and behaviours are however highly individual and the same amount of stress produces different outcomes in both patients and animal models ^8–10^. The resilience as an adaptive response reflecting the ability of the brain to tolerate environmental stress without pathological changes ^11,12^.

Environmental challenges and stress instigate adaptive/maladaptive remodelling of the nervous tissue, which heavily relies on neuroglia, the main homeostatic and defensive arm of the nervous system ^13^. Depression in patients, as well as depression-like behaviours in animal models of major depressive disorder (MDD) and posttraumatic stress disorder (PTSD), are characterised by a prominent decrease in the density and size of astrocytes, which arguably limits the allostatic capacity of the nervous tissue, impairs synaptic transmission, and facilitates the development of aberrant mood ^14–17^. Manipulations with astrocytes and their homeostatic pathways (astrocytes ablation, downregulation of astroglia-specific glutamate transporters or gap junctions) are sufficient to trigger depressive-like behaviours in rodents, whereas manipulating with neurones does not have such an effect ^17–20^. Moreover, stimulating glial glutamate uptake with riluzole or ceftriaxone alleviates depressive-like behaviours and increases astrocytic density ^21,22^. Similarly, treatment with anti-depressive drugs or with acupuncture rescues depressive behaviours and restores astrocytic morphology ^23^. These findings indicate that mood disorders, including stress-induced depression and MDD, are primary astrocytopathies linked to astrocytic atrophy and loss of function.

Astrocytes (which belong to a larger class of astroglia) are principal homeostatic cells of the central nervous system (CNS) supporting nervous tissue at all levels of organisation, from molecular to organ-wide ^24,25^. Protoplasmic astrocytes populating grey matter are characterised by a complex spongiform morphology with several principal processes or branches emanating from the soma; these processes give rise to branches of higher degree and terminal arborisation made from tiny (∼ 100 nm in thickness) leaflets ^26,27^. Morphological plasticity of these leaflets is regulated by the plasmalemmal-cytoskeleton linker ezrin, involved in filopodia formation and astrocyte process motility ^28–30^. Astrocytic leaflets and, less often, primary branches associate with synapses and form the synaptic cradle that fosters and sustains synaptic transmission ^31^. Astrocytes regulate synaptogenesis, support synaptic transmission through numerous transporters that control neurotransmitters, provide ionostasis, supply neurotransmitter precursors and energy substrates, and regulate synaptic extinction ^13,31–35^. Thus astrocytic atrophy limits homeostatic support and hence impairs the synaptic transmission.

Astroglia contribution to the neuropathology is complex and mutable; pathophysiology of astrocytes is ranging from reactive astrogliosis to astrocytopathies, astrocytic atrophy with loss of function, astrocytic degeneration, and astrocytic death ^36^. Astrocytic atrophy is manifest in ageing ^37,38^, neuropsychiatric disorders ^39,40^, neurodegenerative disorders ^41^, epilepsy ^42,43^, aberrant social behaviour ^44^, fear memories ^45^, and addiction ^46^; the loss of astrocyte homeostatic support is one of the leading mechanisms of neuronal damage and death across many neurological diseases ^36^. Retraction of astrocytic leaflets is of a particular importance, as it affects synaptic support and causes aberrant synaptic transmission and neuronal excitability. Chronic unpredictable mild stress (CUMS) is generally employed to generate rodent models of depression ^47,48^. Exposure to various stress regimens, including CUMS, does not trigger depressive-like behaviour in all mice, a sub-population (15 – 50%) of animals demonstrate resilience to stress, which arguably reflects various adaptations ^10,49–52^. Morphological examination of CUMS-induced depressed mice revealed significant atrophy of astrocytes, manifested by a decrease in GFAP-positive profiles as well as astrocytic domains visualised with specific expression of genetic fluorescent probes ^23,53,54^.

In our previous study we found a correlation between astrocytic atrophy, down-regulation of ezrin, and depressive-like behaviours ^23^, suggesting a causal link between ezrin and pathophysiology of depression. In the present study, we manipulated with astrocytic ezrin expression and analysed astrocytic morphology using high-resolution morphological reconstructions and behaviour of mice subjected to CUMS regimen. We found that ezrin overexpression not only prevented astrocytic atrophy but also significantly increased mice resilience to stress. Our work suggests the key role of ezrin and astrocytic leaflets in the pathophysiology of depressive disorders and defines ezrin as a target for managing stress-related depression.

## Results

### Astrocytic ezrin expression defines resilience to chronic stress

To evaluate the role of astrocytic ezrin in mice response to stress we subjected to CUMS protocol three experimental groups: (i) mice which received an injection of a vector with mCherry to label astrocytes (CUMS group; n = 103); (ii) mice with astrocyte-specific overexpression of ezrin (Ezrin-OE; n = 20), and (iii) mice with astrocyte-specific knockdown of ezrin (Ezrin-KD, n = 20). Both overexpression and knock-down were induced by relevant viral-carried constructs, as described in method section (experimental design is shown on Fig. 1a). Virus was stereotaxically injected into right PFC as shown on Fig 1b; which also demonstrates the efficacy of ezrin overexpression and knock-down as revealed by immunocytochemistry. We also set up the control group, which received an injection of a vector with mCherry to label astrocytes but were not exposed to CUMS (Control group, n = 23). After 4 weeks of mice exposure to CUMS behavioural tests and post-mortem analyses were performed. The tests included (i) sucrose intake (indicator of anhedonia), (ii) tail suspension test, TST, (iii) forced swimming test, FST (both measuring the degree of failure of escape-like behaviours, indicative of vulnerability to stress), and (iv) open field test, OFT, a measure of exploratory behaviour and anxiety ^55–57^. We analysed behavioural tests, considering the number of positive tests (i.e. indicative of depressive-like behaviour) for each animal; for behavioural tests thresholds see methods section. As shown on Fig. 1c in the CUMS group majority of animals demonstrated depressive-like behaviours although a sub-population (18%) did not develop such behaviours at all indicating resistance to stress. In the Ezrin-OE mice, in contrast, most of the animals showed resistance to stress with 25% having all tests negative; 40% showing only 1 positive test, 40% - 2, only 16% showing 3 and none showing all 4 tests indicative of depressive like behaviour). Finally, animals form Ezrin-KD group were more sensitive to stress, with 60% of animals showing 3 or 4 positive tests, and 8% 2, and 4% 1. None of Ezrin-KD animals was fully stress-resistant. The sensitivity of mice to CUMS correlated with expression of ezrin in the PFC quantified by immunocytochemistry (Fig. 1d). Thus we concluded that the level of astrocytic ezrin defines mice resilience/sensitivity to chronic stress.

**Figure 1.**
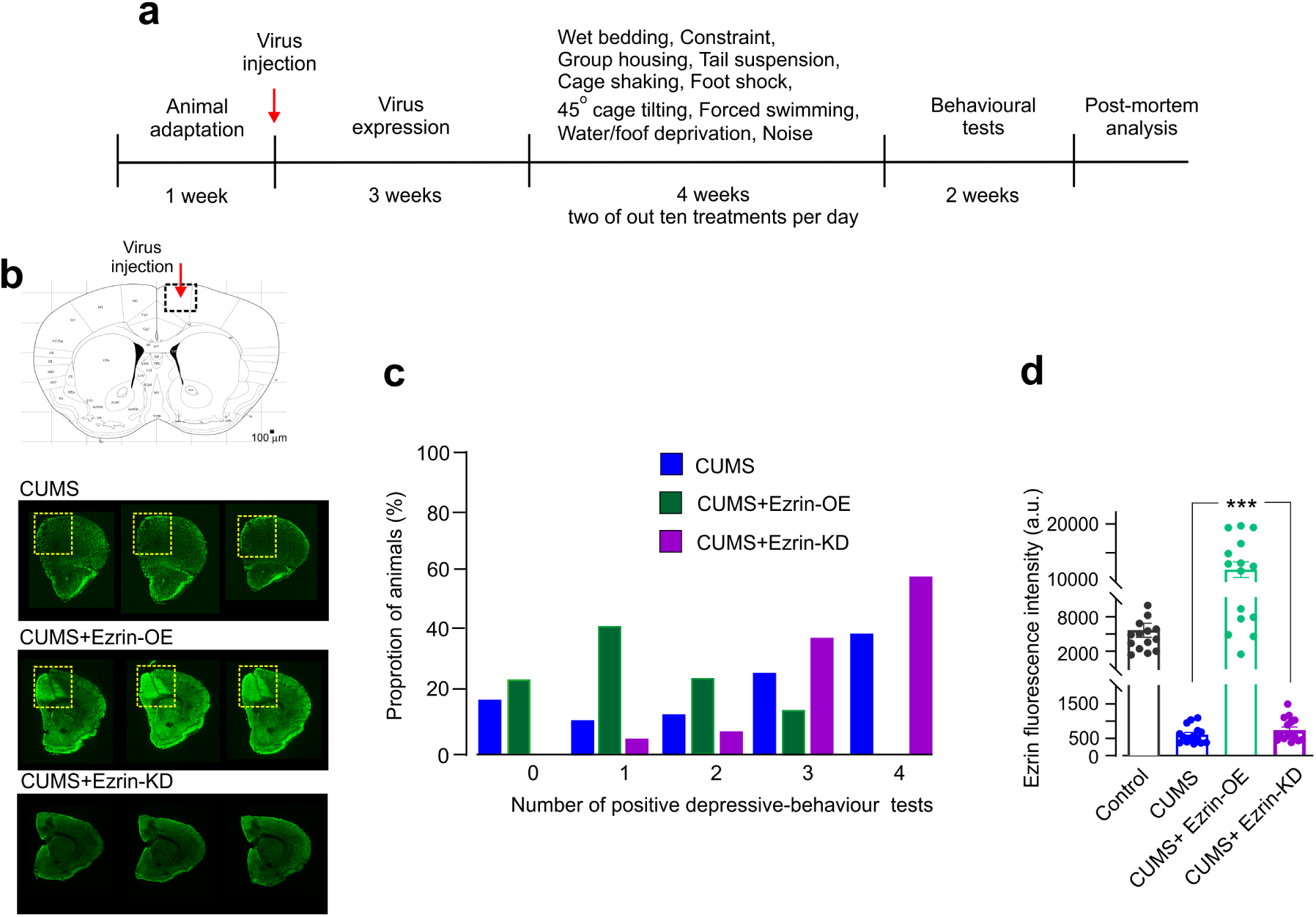
Astrocytic ezrin expression defines mice resilience/sensitivity to susceptibility to chronic stress. a. Experimental protocol. b. The map of mouse brain with the virus injection site in the right prefrontal cortex and immunostaining of the PFC with anti-ezrin antibodies showing efficacy of both overexpression and knock-down of ezrin in astrocytes. c. Proportion of animals (in % of the respective groups) demonstrating 1,2,3, or 4 positive depressive-like behaviours tests following 4 weeks of chronic stress regimen. d. Expression of ezrin as quantified by immunocytochemistry in PFC slices in untreated controls, CUMS, CUMS + Ezrin-OE and CUMS + Ezrin-KD animal groups. ***p < 0.001.

For subsequent astrocyte analysis we selected animals to set up 5 experimental groups of 6 mice each. The first (Control) was composed from 6 mice randomly selected from animals not exposed to CUMS. To the second group (CUMS-sensitive) we assigned (again randomly) mice showing 4 positive tests. To the third group (CUMS-resistant) mice which showed no positive tests were allocated; the fourth group was made from Ezrin-OE animals showing maximal resistance to stress and the fifth group included Ezrin-KD mice with 4 positive tests each.

The average data obtained from behavioural tests for mice groups set up as described above are shown in Fig. 2. The sucrose consumption (defined from consumed sucrose solution as a percentage of total amount of liquid taken in during 12 h that was measured by liquid taken volume), reflecting development of anhedonia, was 87 ± 2% in control, 58 ± 0.8% in CUMS-sensitive mice, (p < 0.001 versus control), 90 ± 2% in CUMS-resistant mice (P < 0.0001 versus CUMS-sensitive), 83 ± 3% in Ezrin-OE group (P < 0.001 versus CUMS-sensitive), and 62% ± 2% for Ezrin-KD (P < 0.0001 versus control) (Fig. 2a).

**Figure 2.**
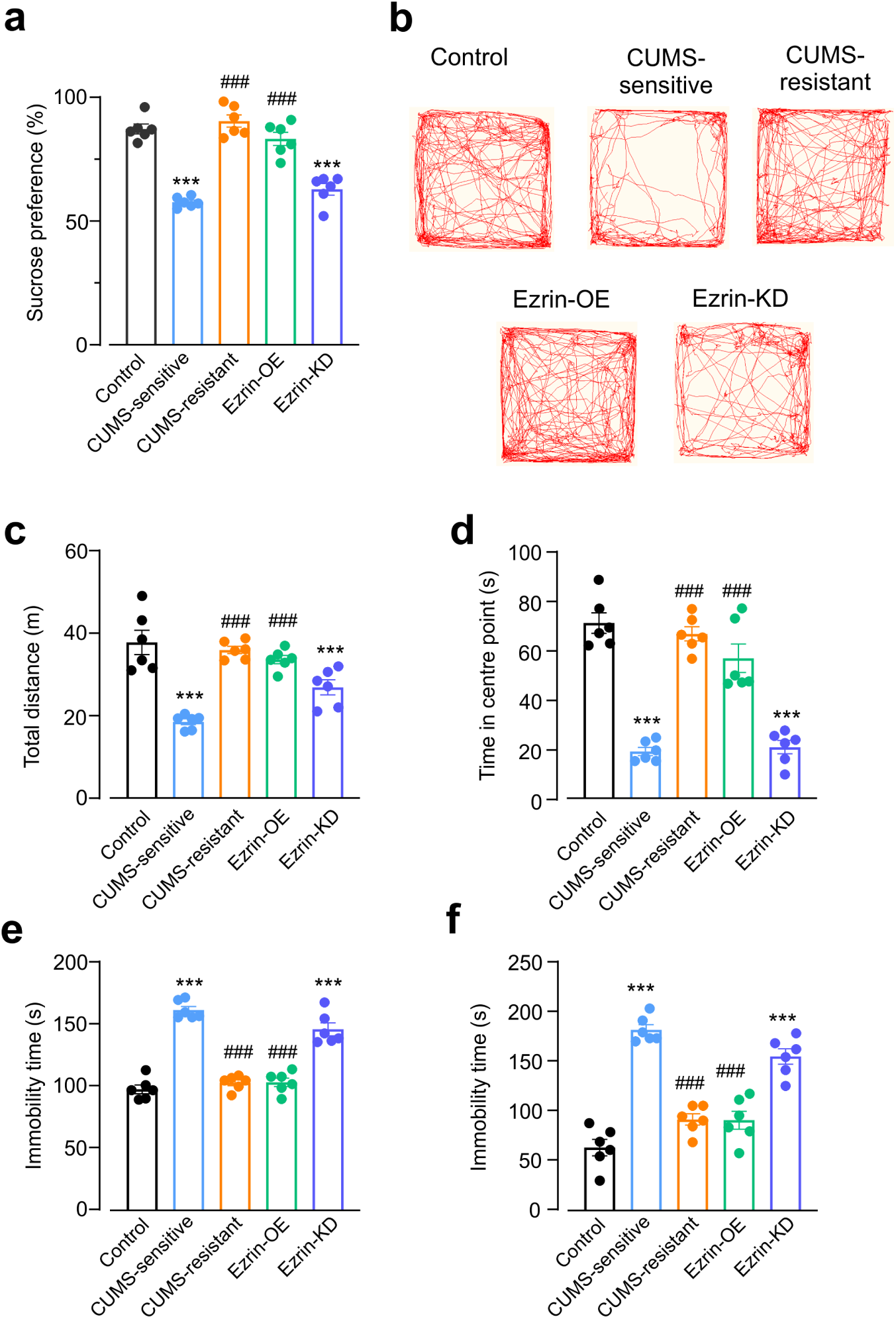
Behavioural phenotypes of mice subjected to CUMS protocol. a. sucrose intake in mice form different experimental groups. b-d. exploratory behaviour in open field test; b. Representative running trace in open field test, the observation time was 10 min. c. Total running distance. d. Centre-point cumulative duration. e,f. Immobility time of mice in tail suspension test and forced swimming test respectively. All data are presented as mean ± sem. *versus control group, # versus CUMS-sensitive group. ***/###p < 0.001.

In the open field test (Fig. 2b,c) the running distance for control mice was 37.74 ± 2.94 m; in CUMS-sensitive mice it was significantly shorter - 21.30 ± 1.01 m (p < 0.0001). In CUMS-resistant group the running distance was 35.89 ± 0.89 m being thus similar to control animals (P = 0.9346), likewise it did not differ significantly from Ezrin-OE group (33.59 ± 1.00 m, P = 0.4237). In Ezrin-KD group the running distance was 26.16 ± 1.56 m (significantly smaller than in control, in CUMS resistant and in Ezrin-OE groups P = 0.005, 0.0032, 0.0327 respectively, and not different from CUMS-sensitive mice; p = 0.2743). Centre point cumulative time was 70.78 ± 4.119 s in control, 18.89 ± 1.680 s in CUMS-sensitive mice (P = <0.0001 compared to control), 66.41 ± 2.894 s in CUMS-resistant mice, 56.55 ± 5.773 s in Ezrin-OE mice, 20.65 ± 2.724 s in Ezrin-KD mice.

Depressive-like behaviours assessed with TST and FST (Fig. 2e, f) showed the following outcomes. In control the TST immobility time was 97.83 ± 3.56 s, in CUMS-sensitive mice 161.8 ± 2.97s (P <0.0001to control), in CUMS-resistant 103.2 ± 2.37s, in Ezrin-OE 103.5 ± 3.44 s (both showing no difference to control P = 0.8345 and 0.8045 respectively, but both being significantly shorter than in CUMS sensitive group P < 0.0001). In Ezrin-KD group the immobility time was 146.3 ± 5.18 s. In FST the immobility time for control group was 63.67 ± 8.31s, in CUMS-sensitive group 182.5 ± 5.28 s (P < 0.0001 to control) in CUMS-resistant mice 92 ± 5.72s, and Ezrin-OE mice 91.33 ± 9.04 s (both showing no significant difference from control, P = 0.0803 and 0.0915 respectively, and both being significantly shorter than CUMS-sensitive group). In Ezrin-KD group the immobility time was 155.7 ± 7.839 s (P < 0.0001 versus control).

### Stress and manipulation with ezrin expression affect astrocytic morphology

First, we analysed morphology of astrocytes using confocal imaging of astrocytes labelled with cell targeted genetic indicators (see methods). Astrocytes from the PFC of mice sensitive to CUMS, show significant morphological atrophy and loss of complexity (Fig. 3a), evidenced by (i) decrease in the length of primary branches (Fig 3b), (ii) decrease in astrocytic territorial domains measured as the area of the maximal projection of an astrocyte along Z-axis (Fig. 3c), and (iii) decrease in the number of intersections in the Sholl analysis (Fig 3d,e). No significant morphological atrophy however was detected in mice resistant to CUMS. The size of territorial domain, branches length and numbers of intersections in control mice were, respectively 660.5 ± 48.31 μm^2^, 13.70 ± 0.49 μm, and 14.80 ± 0.86, in CUMS-sensitive mice these parameters were 359.4 ± 19.88 μm^2^, 10.30 ± 0.23 μm, and 8.5 ± 0.39 (P < 0.0001, to the control) and in CUMS-resistant group they were 574.5±39.54 μm^2^, 12.98 ± 0.40 μm, and 14.75 ± 1.13 (P = 0.4651, > 0.99,> 0.99 to the control). Ezrin overexpression prevented stress-induced astrocytic atrophy whereas Ezrin-KD mice showed astrocytic atrophy similar to that of mice from CUMS-sensitive group (territorial domain, branches length and numbers of intersections were 565.0 ± 42.42 μm^2^, 12.98 ± 0.52 μm, and 14.80 ± 0.60, in Ezrin-OE, and 307.5 ± 32.15 μm^2^, 8.94 ± 0.29 μm, 10.90 ± 0.34 in Ezrin-KD groups respectively, Fig. 3).

**Figure 3.**
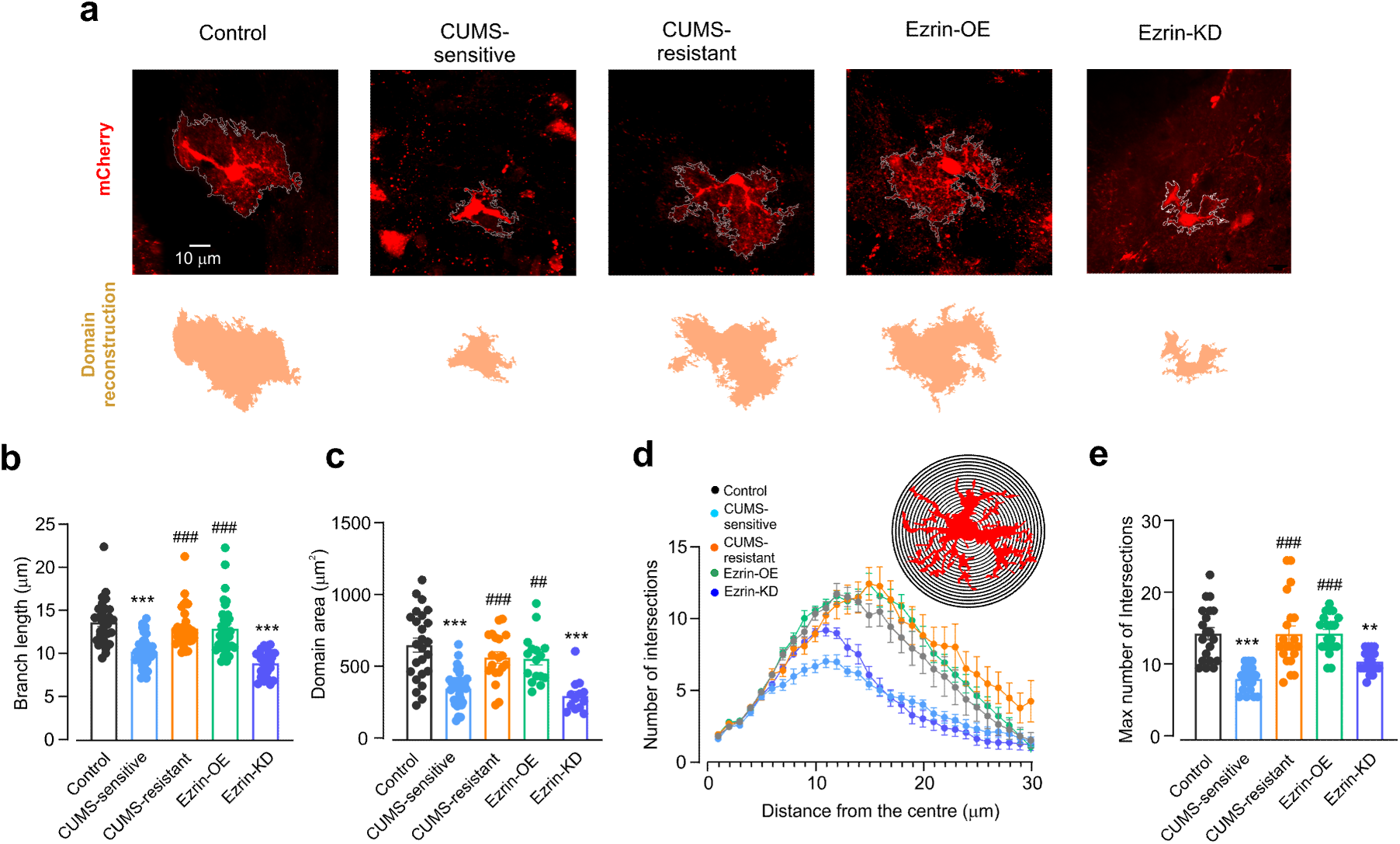
Morphology of PFC astrocytes from CUMS treated animals is affected by intrinsic animal sensitivity to stress and by manipulation with astrocyte ezrin expression. a. Representative confocal astrocytic images and domain area of astrocytic profiles. b. Average branch length of astrocyte, n=25-50 cells, 3 mice c. Astrocytic territorial domains, n=13-35 cells, 3 mice. d. Sholl analysis of astrocytic morphology shows the number of intersections of astrocytic branches with concentric spheres centred in the middle of cell soma, n=20 cells, 3 mice. e. Maximal number of intersections for astrocytes, n=20 cells, 3 mice. All data are presented as mean ± sem. *versus control group, # versus CUMS-sensitive group., **/##p < 0.01, ***/###p < 0.001.

To analyse astrocytic morphology in more details we performed confocal imaging on PFC astrocytes injected with Lucifer yellow in cortical slices (see ^44^ and method section, for the detailed description of the technique). This approach allows much more insight into fine astrocytic morphology as shown on Fig. 4 which demonstrates representative 2D images, territorial domains and 3D reconstructions of astrocytes from different experimental groups. In addition, complexity of astrocytes was assessed by Sholl analysis. Confocal imaging of Lucifer yellow injected astrocytes further corroborated morphological changes induced by stress and manipulation with astrocytic ezrin expression. Astrocytes from CUMS-sensitive mice showed prominent morphological atrophy (Fig. 4a-d); whereas CUMS-resistant astrocytes did not differ form the control group (the size of territorial domain, branches length and numbers of intersections in control mice were, respectively 1470 ± 62.72 μm^2^, 17.64 ± 0.84 μm, and 60.00 ± 3.27, in CUMS-sensitive mice these parameters were 897.4 ± 47.58 μm^2^, 11.37 ± 0.39 μm, and 36.45 ± 1.49, P < 0.0001 to the control; while in CUMS-resistant group they were 1495 ± 69.73 μm^2^, 15.28 ± 0.54 μm, and 62.64 ± 3.83, P > 0.99, > 0.99,= 0.9568 to the control). Exposure to stress did not significantly alter morphology of astrocytes from Ezrin-OE animals, whereas astrocytes from Ezrin-KD group showed significant atrophy (territorial domain, branches length and numbers of intersections were 1338 ± 76.23 μm^2^, 15.09 ± 0.83 μm, and 56.73 ± 2.39, in Ezrin-OE, and 800.3 ± 88.35 μm^2^, 9.553 ± 0.38 μm, and 34.73 ± 1.67 in Ezrin-KD groups respectively). We also quantified the volume fraction (VF ^58^) of astrocytic leaflets (Fig. 4f) to assess changes associated with stress and manipulation with Ezrin expression. The VF of optically unresolved processes was estimated as a ratio of fluorescence along the astrocytic anatomic domain cross-section to the maximal fluorescence of soma as described in ^59,60^. Mean VF was significantly reduced in the astrocytes from CUMS-sensitive and Ezrin-KD mice when compared to control (Control: 11.20 ± 1.00; CUMS-sensitive: 4.06 ± 0.60; Ezrin-KD: 3.67 ± 0.45; P < 0.0001 to control for both), in CUMS-resistant and Ezrin-OE animals VF did not differ significantly from the controls (CUMS-resistant 10.45 ± 0.89, P = 0.9701; Ezrin-OE 10.92 ± 0.97, P = 0.9992).

**Figure 4.**
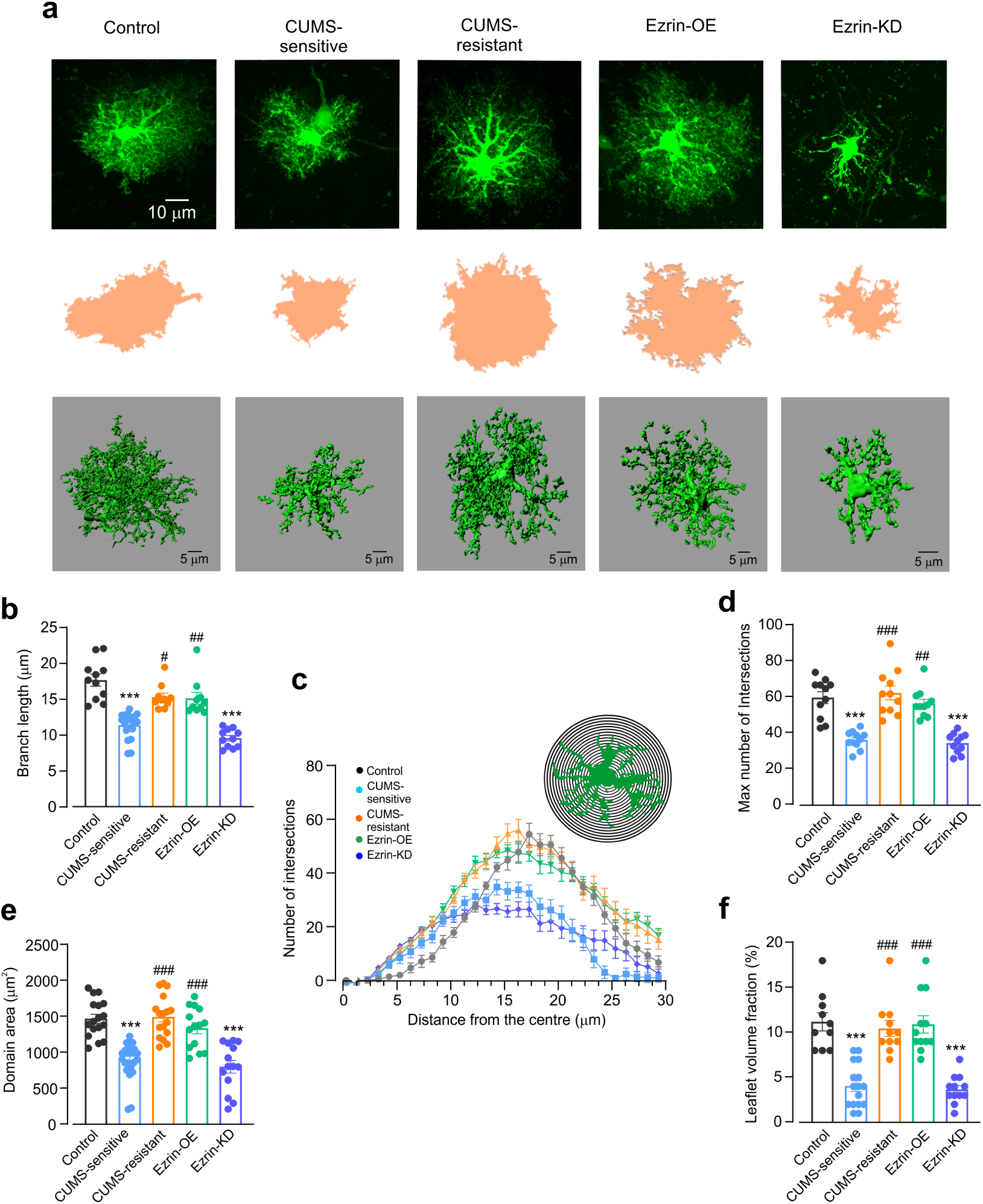
Fine morphology of astrocytes subjected to CUMS a. Representative confocal images (top), domain projection (middle) and 3D reconstruction profile (bottom) of Lucifer yellow labelled astrocytes. b. Average length of astrocytic processes, n=10-20 cells, 3 mice. c. Sholl analysis of astrocytic morphology shows the number of intersections of astrocytic branches with concentric spheres centred in the middle of cell soma, n=11 cells, 3 mice. d. Maximal number of intersections for astrocytes, n=11 cells, 3 mice. e. Astrocytic territorial domains, n=14-26 cells, 3 mice. f. Astrocytic volume fraction, n=10-16 cells, 3 mice. All data are presented as mean ± sem. *versus control group, # versus CUMS-sensitive group. */#p < 0.05, **/##p < 0.01, ***/###p < 0.001.

### Ezrin and phospho-ezrin expression in astrocytes resistant and sensitive to stress

We quantified Ezrin expression with immunoassay against Ezrin and phosphorylated Ezrin (p-ezrin, the active form of this linker) in all groups. Ezrin was revealed as puncta associated with fluorescently labelled astrocytic profiles (Fig. 5a shows representative images of astrocytes, ezrin and p-ezrin immunoreactivity and 3D reconstructions of astrocytes and p-ezrin puncta). Exposure to CUMS led to a significant decrease in both ezrin and p-ezrin in CUMS-sensitive mice; with no changes in these proteins in CUMS-resistant animals. As expected ezrin intensity was high in Ezrin-OE group, and low in Ezrin-KD group. Quantification of expression of ezrin and p-ezrin in all experimental groups are shown in Fig 5b. The colocalisation of ezrin and p-ezrin with soma and processes of astrocytes quantified as a ratio of total ezrin surface area versus total area of astrocytic profile was as follows. Soma: Control group 113.8 ± 15.54; CUMS-sensitive group 44.21 ± 4.57 (P < 0.0001 to control), CUMS-resistant group 88.50 ± 13.29, Ezrin-OE group 74.78 ± 2.89 (P = 0.0241, 0.0120 compared with CUMS-sensitive, respectively), Ezrin-KD 54.54 ± 7.40 (P = 0.0346 compared to control). Association of ezrin with astrocytic arborisation showed similar changes and was 438.1 ± 40.87 for control mice, 73.09 ± 8.17 for CUMS-sensitive, 338.5 ± 48.49 for CUMS-resistant, 266.4 ± 32.17 for Ezrin-OE, 115.9 ± 18.95 for Ezrin-KD (Fig. 5b).

**Figure 5.**
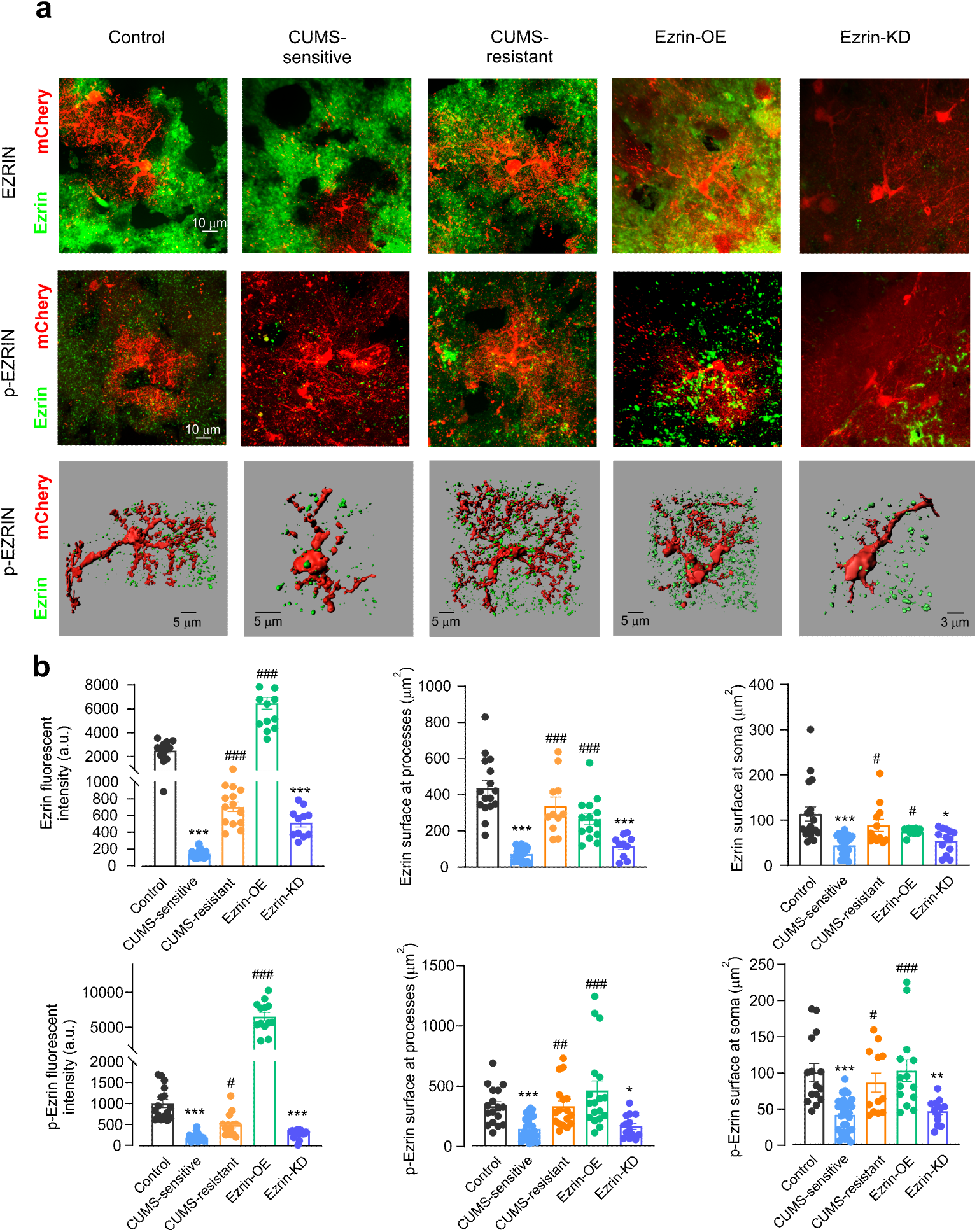
Astrocytic morphology and ezrin expression: direct correlation. a. Representative confocal astrocytic images with Ezrin staining (top), p-Ezrin staining (middle); representative 3D-reconstruction (bottom) of astrocytic profiles (red) with p-Ezrin puncta (green). b. Average fluorescence intensity of Ezrin and p-Ezrin, n=11-24 cells, n=13-32 cells, 3 mice (left column) and surface area of Ezrin and p-Ezrin puncta associated with processes, n=10-22 cells, n=14-30 cells, 3 mice, (middle column) and soma of 3D reconstructed astrocytes, n=9-24 cells, n=12-33 cells, 3 mice (right column). All data are presented as mean ± sem. *versus control group, # versus CUMS-sensitive group. */#p < 0.05, **/##p < 0.01, ***/###p < 0.001.

p-Ezrin puncta surface to astrocytic soma surface colocalisation was significantly higher than in CUMS-sensitive group (CUMS-resistant: 85.25 ± 13.09 versus 40.71 ± 4.05 in CUMS, P = 0.0422; Ezrin-OE: 101.8 ± 14.99 versus. 40.71 ± 4.05 in CUMS, P = 0.0002, Fig. 5b). Colocalisation of p-Ezrin with astrocytic branches also showed same trend when compared with CUMS-sensitive group (CUMS-resistant: 327.6 ± 44.99 versus 140.8 ±16.89, P = 0.0016; Ezrin-OE: 458.2 ± 79.38 versus 140.8 ± 16.89,P < 0.0001 (Fig. 5b).

## Discussion

Astrocytes are fundamental component of the neural network in the CNS, being indispensable for nervous tissue homeostasis and defence ^36,61,62^. In particular, astrocytes regulate neurotransmission and support synaptic connectivity through multiple mechanisms including regulation of synaptogenesis, control of synoptic maturation, support of homeostasis of synaptic cleft and catabolism of major neurotransmitters, as well as overseeing synaptic elimination ^31,34,63,64^. Deficient astrocytic homeostatic support and failed neuroprotection are fundamental pathophysiological mechanisms that contribute to pathogenesis of numerous neurological and neuropsychiatric diseases ^36^. Astrocyte-synaptic interactions are of particular relevance for regulation of neurotransmission in both physiological and pathological contexts. These interactions are mainly centres at astrocytic leaflets that establish contacts with synapses thus contributing to a multicomponent synaptic units which integrate pre- and post-synaptic neuronal compartments with processes of astrocytes, microglia and oligodendrocyte precursor cells ^26,31,65–67^. Morphological plasticity of astrocytic leaflets defines the presence of the latter within the synaptic landscape and modulates synaptic transmission ^68^. Retraction or extension of leaflets significantly affects synaptic kinetics of glutamate, glutamate spillover and K^+^ buffering thus modulating the course of synaptic transmission and modulating synaptic plasticity ^60,69^. In physiological settings astrocytic morphological changes are transient and reversible, in pathology they became rigid and long-lasting thus, arguably, contributing to aberrant synaptic transmission intimately involved in pathophysiology of various psychiatric diseases including disorders of mood.

Astrocytic atrophy in depressive disorders and in stress-induced depressive behaviours is well documented ^17,53,54^. Furthermore, causal relations between morphological presence and functional activity of astrocytes and behavioural changes are widely recognised, whereas anti-depressant therapies restore astrocytic morphology together with rescuing behavioural deficits ^18,21,23^. In the present study we revealed the plasmalemma-cytoskeletal linker ezrin as a key molecule responsible for astrocytic morphological remodelling in response to chronic stress, and moreover we found that manipulating with ezrin expression in astrocytes from PFC affects the susceptibility of mice to stress challenge. We demonstrated that astrocyte-specific ezrin overexpression markedly increases mice resistance to stress, and vice versa, down-regulation of astrocytic expression of ezrin facilitates the development of depressive-like behaviour following exposure to CUMS. In parallel, ezrin expression correlates with (and arguably defines) astrocytic morphology. Mice responses to the chronic stress are heterogeneous and not every animal develops a depressive-like behaviours; this has been well established in animal models and reflects human pathology ^10,12^. The stress susceptibility and resilience are characteristic feature of depressive disorders in patients, the balance between the two is defined by both anatomical and functional idiosyncrasies of every given individual ^9,11^. In this respect stress-resilience closely borders the cognitive reserve, which defines cognitive outcome of various neurological diseases ^70,71^. The molecular and anatomical background for stress resilience remains largely unexplored. We hypothesized that changes in expression of plasmalemmal-cytoskeletal linker ezrin, which belongs to ezrin-radixin-moesin family and is expressed predominantly in astrocytic leaflets ^30^ contributes to stress-induced atrophy of astrocytes.

We characterised in depth relations between astrocytic morphology and ezrin expression in the context of susceptibility to stress. To this end, we segregated the naïve mice into those showing full resistance to stress from those showing maximal sensitivity to it and analysed them separately. We found that neither astrocytic morphology nor ezrin presence in astrocytes in mice resilient to stress differed substantially from control animals not subjected to stress. Similarly astrocytic morphology was not changed in mice over-expressing ezrin and subjected to CUMS protocol. To the contrary, astrocytes demonstrated profound atrophy and decrease in ezrin expression in CUMS-sensitive naïve animals and in Ezrin-KD mice; all major morphometric parameters such as complexity, size of territorial domain, and branch length were significantly decreased. Astrocytic atrophy was analysed in more details by imaging astrocytes injected with Lucifer yellow – these images confirmed profound atrophy of astrocytes and revealed significant decrease in volume fraction occupied by astrocyte leaflets relevant for synaptic coverage. This atrophy was evident in CUMS-sensitive and Ezrin-KD animals; neither CUMS-resistant nor Ezrin-OE mice showed signs of atrophy or decrease in volume fraction of leaflets.

How can astrocytic atrophy and withdrawal of leaflets contribute to the pathophysiology of major depression? The answer probably lies in the substantial remodelling of the synaptic landscape leading to aberrant neurotransmission. At the same time retraction of leaflets from synapses modifies glutamate clearance and K^+^ buffering both contributing to changes in neuronal excitability, information processing and memory ^45^. In particular, depressive-like behaviours in animals and depression in patients are associated with failure in glutamate homeostasis ^50,72^. In particular an increased serum glutamate was detected in patients with MDD ^73^, and moreover the levels of serum glutamate positive correlated with the severity of depression ^73^. Aberrant glutamate homeostasis may also translate into loss of spines and dendritic shrinkage in hippocampal neurones as well as in decreased neurogenesis ^74^. Astrocytes are central for regulation of glutamate levels in the interstitium through the glutamate uptake mediated by excitatory amino acid transporters EAAT1 and EAAT2 ^75^. Retraction of astrocytic leaflets, where majority of glutamate transporters are concentrated ^76^, from the synapses compromises glutamate clearance thus increasing glutamate levels and affecting glutamate homeostasis. Furthermore reduced astrocytic homeostatic support may directly affect neuronal adaptation and alter neuronal ensamples, which in turn are responsible for behavioural changes. Preventing astrocyte atrophy through cell-specific expression of ezrin increases stress resilience and reduces emergence of depressive-like behaviours. Thus ezrin acts a molecular target that translates stress into behavioural outcome through affecting astrocytes, which in turn precipitate changes in neuronal architecture and aberrant mood. How indeed stress acts on astrocytes and what is the molecular link? This requires further investigations. Stress most likely acts on the brain through activation of hypothalamic-pituitary-adrenal axis that affects various aspects of nervous system including metabolism, hormonal control, excitability, and ultimately function ^77–79^. How are astrocytes implicated and what is the mechanistic link to astrocytic morphology? The latter is under noradrenergic control ^80^, which may be affected by activation of HPA. Future focused studies are needed to resolve this matter.

## Conclusion

In summary, we demonstrated that astrocytic expression of ezrin is directly linked to astrocytic atrophy and consequently to the susceptibility of animals to stress. Increase in ezrin expression prevents astrocytic atrophy and favours stress-resilience; to the contrary decreased expression of ezrin promotes atrophy and facilitates development of depressive-like behaviours in response to stress. Our study provides experimental evidence supporting an idea that Ezrin boosts astrocytic presence in the brain active milieu thus protecting it against stress-induced pathological remodelling resulting in mood disorders.

## Materials and Methods

### Animals

All experiments were performed on C57BL/6 mice (obtained from Chengdu Dossy Experimental Animal Co., Chengdu, China); the mice 6 weeks old at the beginning of the experimental protocol, which lasted 10 weeks (Fig. 1a). All mice were adapted to the standard laboratory conditions (24 ± 2 °C room temperature and 65 ± 5% humidity on 12/12 h light-dark cycles) with drinking water and food available *ad libitum*. No statistical methods were used to pre-determine sample sizes, but our sample sizes are similar to those reported in previous publications ^48,54^. The experimental procedures were made in accordance with the National Institute of Health Guidelines for the Care and Use of Laboratory Animals and approved by the Animal Ethics Committee of Chengdu University of Traditional Chinese Medicine (protocol code, AM3520, 8 May 2019).

### Experimental groups

We established five experimental animal groups (to which mice were randomly assigned) before behavioural tests: (i) control group; mice were injected with mCherry construct to label astrocytes, and were not exposed to CUMS; (ii) CUMS group - animals were injected with mCherry construct and were exposed to CUMS; (iii) Ezrin-OE Ezrin-overexpression group - animals exposed to CUMS and injected with both mCherry and Ezrin-overexpression virus; (iv) Ezrin-KD group, animals exposed to CUMS and injected with blue fluorescent protein and Ezrin-knock down virus.

### Chronic unpredictable mild stress (CUMS) regimen

Mice were exposed to the random sequence of stressors during each 24-h period for 4 weeks, as previously described ^23,81^. These stressors included water and food deprivation (12 h), cage tilt 45° (12 h), group housing (12 h), swimming in 4 °C water (5 min), foot shock (1 mA, 5 min), noise (120 dB for 3 h), tail suspension (5 min), damp bedding (12 h), cage shaking (40/min for 5 min), and restraint (1 h).

### Behavioural tests

#### Sucrose preference test

The sucrose preference test is a reward-based test and a measure of anhedonia, and it was performed as previously described ^82^. The mice were singly caged for 3 days and given two 50 mL bottles containing water or water-based 1% sucrose solution (wt/vol), respectively. The bottle positions were switched daily to avoid a side bias. Following a 24 h period of water and food deprivation, the preference for sucrose or water was determined overnight. Sucrose preference (%) was quantified as (volume sucrose/(volume sucrose+ volume water)) × 100%.

#### Tail suspension test

The tail suspension test is a behavioural despair based test assessing the duration of immobility of mice subjected to inexorable conditions, as previously described ^83^. Each mouse was suspended by its tail at a height of 20–25 cm by using a piece of adhesive tape wrapped around the tail 1 cm from the tip. Behaviour was recorded for 6 min. The time of immobility was analyzed during the last 4 min of the 6-min testing period, which followed 2 min of habituation. The duration of immobility was calculated by an observer blinded to the treatment groups. The mice were considered to be immobile only when they remained completely motionless; mice that climbed along their tails were not included.

#### Forced swimming test

The FST was performed as previously described ^84^, in a clear glass cylinder filled with water (temperature, 23–25 °C); cylinder’s dimensions were: height, 30 cm; diameter, 20 cm; water level, 15 cm. Mice were gently placed in the tanks. Following the swimming session, the mice were removed from the water by their tails, gently dried with towels, and kept warm under a lamp in their home cages. They were considered to be immobile whenever they stopped swimming and remained floating passively, still keeping their heads above the surface of the water. The time of immobility was analysed during the last 4 min of the 6-min testing period, which followed 2 min of habituation ^85^

#### Open field test

The open field test was performed as previously described ^23^. The apparatus consisted of a rectangular chamber (40 × 40 × 40 cm) made of white, high-density, non-porous plastic. Mice were gently placed in the centre of the chamber and their motility was recorded for 10 min. The total running distance, and the time spent in the centre versus the periphery of the open field chamber were recorded by a camera connected to a computer using an automated video tracking program (EthoVision XT 9.0; Noldus, Wageningen, The Netherlands). The chamber was thoroughly cleaned with 95% ethanol and double distilled water, and dried prior to use and before subsequent tests, to remove any scent clues left by the previous subject.

### Behavioural tests threshold

Tail suspension test: immobility time of mice showed more than 120 s within 4 min were considered as depressive-like behaviour.

Force swimming test: immobility time of mice showed more than 100 s within 4 min were considered as depressive-like behaviour.

Sucrose preference test: sucrose preference showed less than 75% were considered as depressive-like behaviour.

Open filed test: total distance less than 35 m and centre-point cumulative time showed less than 30 s were considered as depressive-like behaviour.

### AAVs microinjections

Viral injections were performed 3 weeks before CUMS treatment as indicated in Fig. 1 by using a stereotaxic apparatus (RWD, Shenzhen, China) to guide the placement of a Hamilton syringe fixed with bevelled glass pipettes (Sutter Instrument, 1.0-mm outer diameter) into the PFC ^86^. The injection site was located at half of the distance along a line defined between each eye and the lambda intersection of the skull. The needle was held perpendicular to the skull surface during insertion to a depth of approximately 0.2 mm. A total of 0.7 μl of AAV2/8-gfaABC1D-Ezrin-HAx2-P2A-mCherry-WPRE-pA, AAV5-gfaABC1D-mCherry-WPRE-pA (1 × 10^12^ gc/mL; Taitool Bioscience, Shanghai, China), AAV2/8-gfaABC1D-miRNAi (Ezrin)-BFP-WPRE-SV40,AAV2/8-gfaABC1D-miRNAi(NC)-BFP-WPRE-bGHpolyA(1 × 10^12^ gc/mL; VectorBuilder, Guangzhou, China) was slowly injected into right side of the mPFC. Glass pipettes were left in place for at least 5 min. After injection, animals were allowed to completely recover under a warming blanket and then returned to the home cage.

### Immunohistochemistry

Mice were perfused with cold paraformaldehyde (PFA, 4% w/v in phosphate buffer saline (PBS) under deep isoflurane (2%, 5 min) and pentobarbitone (1%, 50 mg/kg) anaesthesia. Brains were collected, postfixed and cryoprotected in 30% (w/v) sucrose solution. Brains were cut using a cryostat in 45 um thick sections; slices were immediately transferred into storing solution (30% w/v sucrose and 30% ethylene glycol in PBS) and kept at 80 °C until use. Free-floating sections were incubated 1 h in saturation solution (6% fetal calf serum in PBS). The sections were then incubated overnight in the same solution complemented with the primary antibody (rabbit anti-Ezrin 1:100; rabbit anti-p-Ezrin 1:100, Cell Signalling, Danvers, Massachusetts, USA). After washing in PBST three times, slices were incubated 1 h at 37 °C in saturation solution containing the relevant secondary antibody (goat anti-rabbit Alexa 488; Invitrogen, Carlsbad, California, USA). After washing in PBST three times, the coverslips were mounted on slides using anti-fade solution (Solarbio, Beijing, China). Confocal microscopy (Olympus, Tokyo, Japan) or normal fluorescence microscope (Leica, Wetzlar, Germany) were used to obtain images.

### Lucifer yellow injection

Intracellular dye filling was performed on the lightly fixed brain slices from the mPFC of mice using the protocols as previously reported ^87^. In the experiments, mCherry-positive astrocytes in the formalin-fixed brain slice that contained mPFC were visualised. 1.5% Lucifer yellow (Lucifer yellow, Merk, Darmstadt, Germany, #67769-47-5) was iontophoresed into astrocytes. After injection, time-lapse imaging (0.1 Hz) was conducted using a confocal microscope (Nikon A1R+, Tokyo, Japan) with a 40× water-immersion objective lens (numerical aperture, NA 0.8) for 5 min. Finally, the brain slices were mounted on a glass slide by using the anti-fade mounting medium andthe confocal imaging stacks were collected with a Z-step size of 0.25 μm under confocal microscopy (Olympus, Tokyo, Japan).

### Sholl analysis

Sholl analysis is a commonly used method to quantify astrocyte process complexity ^53,88^. All processing steps were performed using image analysis software ImageJ [https://imagej.net/imagej-wiki-static/ Sholl_Analysis]. In brief, Z-stacks corresponding to the emission spectrum (565–610 nm) of mCherry-labelling (resolution was 512 × 512 pixels (0.25 μm/px) on XY axis with a step on Z-axis 1 μm/frame were resampled to the same lateral resolution of 0.25 μm/px.

### 3D reconstructions

The confocal imaging stacks were collected with a Z-step size of 0.25 μm under a confocal microscope (Olympus, Tokyo, Japan). Three-dimensional reconstructions were processed offline using Imaris 7.4.2 (Bitplane, South Windsor, CT) as reported previously ^44,89^. In brief, the astrocyte soma and processes were measured and reconstructed according to their own parameter. Processes diameter was measured as one-tenth of astrocyte soma. In addition, Ezrin, p-Ezrin was measured as 1mm in every group. The surface–surface colocalisation was calculated by an Imaris plugin ^44^.

### Statistics

All statistical analyses were performed by GraphPad Prism 8. All data were expressed as means ± SEM of n observations, where n means the number of animals in behavioural tests, or astrocyte cells from at least three animals. All analysis was single-blind. Data with more than two groups were tested for significance using one-way ANOVA test followed by the Holm–Sidak test. Multiple comparisons between the data were performed in case of their non-normal distribution, using the Kruskal–Wallis ANOVA on ranks, followed by Tukey’s test. A two-way ANOVA followed by Dunn’s test was performed to compare data obtained in Figs. 3d, 4c. Significance was defined as P < 0.05.

## Acknowledgements

This work was supported by grants from NSFC-RSF (82261138557), Sichuan Provincial Administration of Traditional Chinese Medicine (2023zd024), and NSFC (82274668, 82230127).

## Author contributions

SL, AV, YT conceived the study, SL and AV designed experiments, SL, SLL, XYR, JG and TJL performed experiments, BL and BJC provided expertise in CUMS, BZ and RTJ contributed to lucifer yellow experiments and image analysis, AS and PI contributed to discussion, AV and SL wrote the manuscript, AV, YT, PI, AS, CY, JN, BL edited the manuscript.

